# Impact and role of hypothalamic corticotropin releasing hormone neurons in withdrawal from chronic alcohol consumption in female and male mice

**DOI:** 10.1101/2023.05.30.542746

**Authors:** Sofia Neira, Sophia Lee, Leslie A. Hassanein, Tori Sides, Shannon L. D’Ambrosio, Kristen M. Boyt, Jaideep S. Bains, Thomas L. Kash

**Affiliations:** Bowles Center for Alcohol Studies, Department of Pharmacology, School of Medicine, University of North Carolina at Chapel Hill, Chapel Hill, North Carolina, USA; Department of Pharmacology, School of Medicine, University of North Carolina at Chapel Hill, Chapel Hill, North Carolina, USA; Hotchkiss Brain Institute and Department of Physiology and Pharmacology, University of Calgary, Calgary, Alberta, Canada

## Abstract

Worldwide, alcohol use and abuse are a leading risk of mortality, causing 5.3% of all deaths (W.H.O., 2022). The endocrine stress system, initiated by the peripheral release of corticotropin releasing hormone (CRH) from primarily glutamatergic neurons in the paraventricular nucleus of the hypothalamus (PVN), is profoundly linked with alcohol use, abuse, and relapse (Blaine & Sinha, 2017). These PVN CRH-releasing (PVN^CRH^) neurons are essential for peripheral and central stress responses (Rasiah et al., 2023), but little is known about how alcohol affects these neurons. Here, we show that two-bottle choice alcohol consumption blunts the endocrine mediated corticosterone response to stress during acute withdrawal in female mice. Conversely, using slice electrophysiology, we demonstrate that acute withdrawal engenders a hyperexcitable phenotype of PVN^CRH^ neurons in females that is accompanied by increased glutamatergic transmission in both male and female mice. Only male mice show a concurrent increase in GABAergic synaptic transmission. We then tested whether chemogenetic inhibition of PVN^CRH^ neurons would restore stress response in female mice with a history of alcohol drinking in the looming disc test, which mimics an approaching predator threat. Accordingly, inhibition of PVN^CRH^ neurons reduced active escape in hM4Di alcohol history mice only. This study indicates that stress responsive PVN^CRH^ neurons in females are particularly affected by a history of alcohol consumption. Interestingly, women have indicated an increase in heavy alcohol use to cope with stress (Rodriguez et al., 2020), perhaps pointing to a potential underlying mechanism in alcohol mediated changes to PVN^CRH^ neurons that alter stress response.

## Introduction

Alcohol use disorders (AUD) account for the majority of substance use disorders worldwide, with approximately 3 million deaths per year attributed to harmful alcohol use (W.H.O., 2022). Alcohol consumption is also increasing, with recent reports from the United States indicating that women especially are engaging in heavy binge drinking behaviors that are linked to relief from COVID-19 associated stressors (Pollard et al., 2020; Rodriguez et al., 2020). Despite the established relationship between alcohol, withdrawal, and stress (Becker, 2012; Blaine & Sinha, 2017; Peltier et al., 2019; Sinha et al., 2009), few studies have investigated alcohol-induced alterations to paraventricular nucleus of the hypothalamus corticotropin releasing hormone (PVN^CRH^) neurons, which initiate the endocrine stress response via the hypothalamic-pituitary-adrenal (HPA) axis (Kudielka & Kirschbaum, 2005; Stephens & Wand, 2012). In particular, little investigation has been done in female subjects and after long-term voluntary alcohol consumption.

There is a blunting of the HPA-axis stress hormone, cortisol (corticosterone in rodents), to acute stress in both male and female AUD or binge drinking withdrawal subjects (Adinoff et al., 2005; Bernardy et al., 1996; Blaine et al., 2019; Errico et al., 1993; Junghanns et al., 2003; Wand & Dobs, 1991). Baseline cortisol levels in the absence of acute stressors are more varied (Adinoff et al., 2005; Gianoulakis et al., 2003; Keedwell et al., 2001; Sinha et al., 2009; Thayer et al., 2006), though this is more difficult to interpret as many of these measures are taken at varied withdrawal timepoints and less controlled locations. The scarce rodent studies on hormone and PVN response in withdrawal from chronic treatments of alcohol have mostly mirrored human studies. For instance, basal corticosterone and ACTH hormone levels vary, with some studies finding increases (Tabakoff et al., 1978), others decreases (Li et al., 2011; Richardson et al., 2008), and the only study we are aware of that included male and female rats finding that only females showed blunted corticosterone levels two months post alcohol treatment (Silva et al., 2009). Regarding the acute stress response, Lee et al., (2000) subjected 24hr withdrawal male rats to acute stressors and found reduced *Fos* expression in the PVN as well as reduced ACTH levels compared to controls.

Despite rodent investigations offering a direct ability to study alterations to PVN^CRH^ neurons themselves, few studies have explored how alcohol alters these neurons, and fewer still have studied how females are affected. Female rodents show a greater activation of the HPA-axis stress system to different stressors, including acute alcohol challenges, and display more active stress responses in behavioral tasks that activate this system (Bangasser et al., 2018; Kudielka & Kirschbaum, 2005; Palanza, 2001; Palanza & Parmigiani, 2017; Rivier, 1999), perhaps suggesting interactions between chronic alcohol and stress response. In fact, females consume high levels of alcohol in a two-bottle choice (2-BC) voluntary alcohol paradigm, and compared to water controls, display greater active escape behaviors during acute withdrawal in the looming disc test that mimics an approaching predator (Neira et al., 2022). PVN^CRH^ neurons are activated prior to escape initiation in this task, and optogenetic inhibition of these neurons reduces escape behavior, indicating that PVN^CRH^ neurons are necessary for active escape responses to the looming disc (Daviu et al., 2020). Additionally, a history of alcohol has been shown to increase glutamatergic synaptic transmission in PVN^CRH^ neurons in male monkeys (Jimenez et al., 2019) and cause biophysical and synaptic alterations to these neurons in protracted abstinence in rats (Marty et al., 2020; Munier et al., 2022). This evidence led to our hypothesis that PVN^CRH^ neurons are more excitable in acute withdrawal after a history of alcohol consumption, resulting in more active stress coping behaviors. Although some studies suggest blunted endocrine stress system function in withdrawal, hormonal output and alterations to PVN^CRH^ neurons might be detangled. In fact, PVN^CRH^ neurons have recently been implicated in various behaviors independent of HPA-axis mediated hormone release (Daviu et al., 2020; Füzesi et al., 2016; Kim et al., 2019; Yuan et al., 2019).

Despite the prevalence and harm stemming from AUD, few treatments exist to mitigate this chronic relapsing disorder. Relapse rates have been associated with stress response systems, and most often, relapse occurs early in abstinence (Blaine & Sinha, 2017; Junghanns et al., 2003; Karim et al., 2010; Wemm et al., 2019). It is therefore imperative to understand physiological alterations in early withdrawal, particularly those associated with stress response. Thus, we used fluorescence in-situ hybridization, *ex-vivo* slice electrophysiology, and chemogenetic *in-vivo* inhibition to examine how a history of intermittent 2-BC alcohol consumption alters PVN^CRH^ neurons during acute withdrawal in female and male mice. We find drastic effects in PVN^CRH^ neuronal modulation by chronic alcohol intake as well as PVN^CRH^ driven changes in stress coping behavior in alcohol history female mice only.

## Methods

### Two Bottle Choice Alcohol Paradigm

At least 4 days before the start of 2-BC, mice were transferred to a temperature controlled 12-hr reverse dark/light cycle vivarium, single-housed and switched to Isopro RMH 3000 chow (LabDiet, St. Louis, MO), which supports high alcohol intake levels (Marshall et al., 2015). Mice were given *ad-libitum* access to water and food. During the 2-BC procedure, alcohol access female and male mice over 8 weeks of age were given access to water and intermittent 24-hr alcohol (20% w/v) sessions with forced 24-48hr withdrawals between each session (i.e. M/W/F - 24hr Alcohol Sessions: T/Th/Sat/Sun - No Alcohol). Alcohol and water bottles were weighed at the start and end of each alcohol session, which began 2-3 hours into the dark light cycle. Averaged drip values from an empty cage were subtracted from all water and alcohol raw values. Water access control mice were matched for age, weight, and litter and received only water throughout the 2-BC procedure. The Institutional Animal Care and Use Committee at UNC Chapel Hill approved all experimental procedures, which were performed in accordance with the NIH Guide for the Care and Use of Laboratory Animals.

### Home-cage Test

At least 24 hours after the final 2-BC alcohol session, mice were brought to the behavioral testing room and allowed to habituate to the room for a minimum of 40 minutes. After habituation, the home-cage with only the bedding and mouse was placed in a ∼60 lux low-light sound attenuated chamber that contained an overhead camera for filming behavior. Rearing, digging, grooming, and locomotion were analyzed for a 15- minute period using DeepLabCut and SimBA mediated automatic analysis as previously discussed (Neira et al., 2022).

### Plasma Corticosterone and In-situ Hybridization

C57BL/6J (Jackson Laboratories, Bar Harbor, ME) mice were exposed to 18 alcohol sessions in a 2-BC procedure. To measure basal corticosterone and *Fos* mRNA levels in PVN^CRH^ neurons during acute withdrawal, 24-36hrs after the final alcohol drinking session, mice (Females: 10 water controls for corticosterone and 9 for in-situ due to technical errors, 10 alcohol access. Males: 10 water controls, 10 alcohol access) were placed in an isoflurane chamber before being transferred out of their animal facility housing room. A separate set of mice (Females: 11 water controls, 12 alcohol access for corticosterone and 13 for in-situ due to technical errors. Males: 12 water controls, 13 alcohol access) were subjected to the home-cage test 24-36hrs after the final alcohol session and immediately placed in an isoflurane chamber to test corticosterone and *Fos* mRNA levels in PVN^CRH^ neurons during acute withdrawal immediately following an acute stressor. Once fully anesthetized, mice underwent rapid decapitation, trunk blood was collected in a 2mL tube for plasma corticosterone analysis and brains were rapidly frozen over aluminum and dry ice and stored in a -80°C freezer for fluorescence in-situ hybridization (FISH) experiments.

#### Plasma Corticosterone

Trunk blood was centrifuged at 4 °C for 10 min at 3000 RCF and 30µL plasma was extracted and stored at -4°C. To measure corticosterone, the DetectX Corticosterone Enzyme Immunoassay Kit (Arbor Assays, Ann Arbor, MI) was used and followed according to manufacturer instructions. Data was analyzed using a Spectra Max Plate Calorimetric 96 well microplate reader and the MyAssays Arbor Assays analyzer.

#### Fluorescence In-situ Hybridization

Brains were sliced into 12 µL coronal sections containing the PVN using a Leica CM3050 S cryostat (Leica Microsystems) and mounted directly on slides. FISH was performed using the Affymetrix ViewRNA 2-plex assay according to the manufacturer’s instructions using probes for *Crh* (category number: 316091) and *Fos* (category number: 316921-C2). A blinded individual visually determined the slice containing the highest density of CRH neurons concentrated in the medial PVN and chose one hemisphere counterbalanced per mouse for analysis. FISH data was analyzed using QuPath version 0.4.3 (Bankhead et al., 2017). Total number of *Crh* and/or *Fos* positive neurons were counted, and PVN^CRH^ activation was determined by the percent of *Crh* expressing neurons that also expressed *Fos*.

### Ex-vivo Slice Electrophysiology

C-fos is an indirect measure of neuronal activation. Therefore, to more directly test how PVN^CRH^ neurons are affected in acute withdrawal by chronic alcohol consumption, we performed slice electrophysiology using a CRH-reporter mouse strain to visually identify fluorescent, CRH-containing neurons in the PVN. These mice were bred in-house by crossing CRH-ires-Cre (B6(Cg)-Crh^tm1(cre)Zjh^/J) mice to Ai9 cre-dependent tdTomato fluorescent reporter (B6.Cg-Gt(ROSA)26Sor^tm9(CAG-tdTomato)Hze^/J) mice. Mice were exposed to 18-26 sessions of intermittent alcohol using the 2-BC choice procedure and underwent the home-cage test 24-30 hours after the final session. Immediately after, mice were placed in an isoflurane chamber until fully anesthetized and brains were rapidly removed and submerged in an ice-cold carbogen (95% O_2_/5% CO_2_) saturated sucrose artificial cerebrospinal fluid (aCSF) cutting solution (In mM: 194 sucrose, 20 NaCl, 4.4 KCl, 2 CaCl_2_, 1 MgCl_2_, 1.2 NaH_2_PO_4_, 10 D-glucose and 26 NaHCO_3_). Brains were then blocked for the PVN and 250 µm coronal slices were prepared on a Leica VT 1000S vibratome (Leica Biosystems, Buffalo Grove, IL) and transferred to a holding chamber with 34°C heated and carbogen saturated aCSF (in mM: 124 NaCl, 4.4 KCl, 1 NaH_2_PO_4_, 1.2 MgSO_4_, 10 D-glucose, 2 CaCl_2_, and 26 NaHCO_3_). Slices were allowed to equilibrate for at least 1 hour, then transferred to a recording chamber (Warner Instruments, Hamden, CT) with oxygenated and 30±2°C heated flowing aCSF (2mL/min). Neurons were visualized using a 40X immersion objective on a Scientifica Slicescope II (East Sussex, UK) with differential interference contrast and a 550 LED was used to visualize fluorescent, CRH-containing PVN^CRH^ neurons. Whole cell patch-clamp experimental recording signals were acquired using an Axon MultiClamp 700B (Molecular Devices, Sunnyvale, CA), the data was sampled at 10kHz, low-pass filtered at 3kHz, and data was analyzed in pClamp 10.7 (Molecular Devices, LLC, San Jose, CA) or Easy Electrophysiology (Easy Electrophysiology Ltd., UK). One to three neurons were recorded per mouse for each experiment using borosilicate glass capillary micropipettes pulled with a 2-5MΩ electrode tip resistance using a Flaming/Brown P97 electrode puller (Sutter Instruments, Novato, CA). Any changes greater than 20% from the initial access resistance throughout the experiment or a membrane capacitance greater than 30 MΩ led to the exclusion of that neuron from analysis. Cells were also excluded from analysis if the current to hold the cell membrane voltage at -80 mV in current clamp mode exceeded -60 pA.

#### Intrinsic Excitability Experiments

Experiments were recorded in current-clamp mode with a potassium gluconate-based intracellular solution (in mM: 135 K-gluconate, 5 NaCl, 2 MgCl_2_, 10 HEPES, 0.6 EGTA, 4 Na_2_ATP, 0.4 Na_2_GTP, pH 7.3, 289–292mOsm). Input resistance was calculated using steps from -70 to -80mV in the current-voltage relationship in voltage clamp immediately after breaking into the cell. Resting membrane potential (RMP) was measured after stabilization in current clamp. Current was injected to hold cells at a common membrane potential of -80mV, and changes in excitability were measured by the frequency of action potentials fired at increasing 10pA current steps (−40 to +80) lasting 250ms. In total, 11 female water, 9 female alcohol, 10 male water, and 9 male alcohol mice were used.

#### Synaptic Transmission Experiments

Excitatory post synaptic currents (EPSCs) were recorded in voltage-clamp mode with a potassium gluconate-based intracellular solution. Spontaneous transmission (sEPSCs total mice: 8 male and female water mice and 7 male and female alcohol-exposed mice) was recorded at -80mV in the presence of 100 µM GABA-A receptor antagonist picrotoxin and miniature events (mEPSCs total mice: 4 water and alcohol females, 4 water males and 6 alcohol males) were recorded in the presence of 100 µM picrotoxin and 500nM sodium channel blocker TTX.

Inhibitory post synaptic currents (IPSCs) were recorded in voltage-clamp mode with a potassium-chloride gluconate-based intracellular solution (in mM: 80 KCl, 70 K-gluconate, 10 HEPES, 1 EGTA, 4 Na2ATP, 0.4 Na2GTP, pH 7.2, 285–290 mosmol).

Spontaneous transmission (sIPSCs total mice: 8 male, 6 female water mice and 8 male and 7 female alcohol-exposed mice) was recorded at -80mV in the presence of 10 µM AMPA and Kainate receptor antagonist DNQX and miniature events (mIPSCs total mice: 4 water and alcohol females, 4 water males and 6 alcohol males) were recorded in the presence of 10 µM DNQX and 500nM TTX.

### hM4Di Chemogenetic Inhibition

CRH^cre^ in-house bred mice aged 6 weeks or older underwent stereotaxic surgery for infusion of mCherry control or hM4Di virus into the PVN. After at least five days of recovery, mice began a 6-week 2-BC procedure. In the final 3 weeks of 2-BC, including the day before the first CNO experiment, mice were subjected to at least 4 saline habituation injections on alcohol off days. 0.5mg/kg i.p. CNO was administered to mCherry control and hM4Di mice for all behavioral tests 30-50 minutes prior to the test. One day after the final, 18^th^, 2-BC alcohol session ended, mice were subjected to the home-cage test immediately followed by a free social interaction test. Three and four days after the final 2-BC session, mice were once again brought to the behavior room, allowed to habituate for at least 40-minutes, and then given a 20-minute exposure to the looming disc arena with a saline injection given 20-30 minutes prior to the exposure. On the 5^th^ day following 2-BC, mice underwent the looming disc test after at least 40-minute habituation to the room. Mice were perfused and unilateral or bilateral PVN injections were verified, as we saw no differences in behavior between unilateral and bilateral hits. After verification, total Ns include - Females: 13 Water mCherry, 13 Alcohol mCherry, 9 Water hM4Di, 11 Alcohol hM4Di. Males: 10 Water mCherry, 10 Alcohol mCherry, 10 Water hM4Di, 12 Alcohol hM4Di.

#### Stereotaxic Surgery

After 4% isoflurane anesthesia induction, mice were transferred to a stereotaxic frame and kept at 1-2% maintenance in oxygen (1-2L/min). The scalp was sterilized (70% ethanol and betadine) and the skull exposed. Two small holes were made in the skull above the injection sites and 200nl (100nl/min) of AAV8-hSyn-DIO-mCherry or AAV8- hSyn-DIO-hm4Di-mCherry virus (Addgene, Watertown, MA) was infused in both hemispheres of the PVN using a 1µL Hamilton syringe (0° angle, mm relative to bregma: AP: - 0.55, ML: ± 0.20, DV: - 5.15). The virus was allowed to diffuse for 5 minutes following infusion.

#### Free Social Interaction

Immediately following the home-cage test as described above, mice were transferred to a fresh cage with bedding and allowed to habituate for 2-minutes. Following habituation, a juvenile, same-sex mouse was placed in the cage and their interactions were filmed for 10 minutes.

#### Looming Disc

Mice had a minimum 3-minute habituation to the looming disc arena (41 x 19 x 20.5 cm mirrored floor plastic arena with a 13 x 12 x 10cm protective shelter in one corner). Following habituation, 5 episodes of the looming, advancing disc paradigm, separated by at least 1-minute between each episode, were triggered when the mouse was not in the shelter or heading towards it. Each episode requires eight seconds, starting with a three second 2cm black disc on a light grey background which then expands to its full 20cm size over 2 seconds and remains on the screen for an additional three seconds. A individual blinded to the experimental manipulation determined whether the mouse escaped to the hut within the 8 second loom presentation. If escape did not occur, the mouse was scored as freezing, or neither escape/freezing for each of the five looms. Freezing was determined as no movement other than breathing for at least 1 second throughout the loom presentation and escape was counted if the mouse entered the shelter at any moment during loom presentation.

### Statistics

Female and male data was analyzed separately due to existing differences in total alcohol intake during 2-BC as shown in Table 1. FISH and corticosterone data were analyzed using unpaired t-tests and Mann-Whitney two-tailed test for water vs alcohol conditions. Slice electrophysiology data was analyzed using two-way repeated measures ANOVA for current, and condition (water or alcohol) and unpaired two-tailed t-tests when only comparing water vs alcohol groups. Behavioral data for the home-cage and social interaction tests in the hM4Di chemogenetic experiment was analyzed using two-way ANOVAs for condition (water vs. alcohol) and virus (mCherry vs hM4Di). Looming disc data was analyzed using Chi-square tests, as the data was not numerical (escape, freeze, or neither escape nor freeze responses). A p-value less than 0.05 was determined to be significant, and Šidák corrected multiple comparisons were conducted when an ANOVA indicated an interaction effect. Statistical tests were conducted in Graphpad Prism 9.2.0 (GraphPad Software Inc, San Diego, CA). ROUT Outlier tests were conducted on all data. One female alcohol mouse was removed from the plasma corticosterone analysis and the home-cage grooming data analysis per the ROUT outlier analysis after they were confirmed as outliers.

**Table 1:** 2-BC first, last and total alcohol intake per experiment in g/kg. Data shown as average ± standard deviation. Total alcohol intake female and male data analyzed using unpaired students t-test. a** p <.01; b, c, d, and e**** p<.0001.

| Experiment: | Sex: | First session intake (g/kg) | Last session intake (g/kg) | Total intake (g/kg) |
| --- | --- | --- | --- | --- |
| Baseline FISH and Corticosterone | Females: | 15.04 ± 10.28 | 24.53 ± 6.78 | 420.06 ± 107.24<sup>a</sup> |
|  | Males: | 20.13 ± 5.58 | 16.04 ± 2.53 | 300.00 ± 40.60<sup>a</sup> |
| Acute Stress FISH and Corticosterone | Females: | 25.00 ± 9.42 | 29.57 ± 5.82 | 502.19 ± 60.58<sup>b</sup> |
|  | Males: | 20.64 ± 5.23 | 17.95 ± 4.84 | 343.66 ± 72.38<sup>b</sup> |
| Slice Electrophysiology | Females: | 20.57 ± 6.85 | 29.19 ± 6.96 | 543.62 ± 130.25<sup>c</sup> |
|  | Males: | 16.12 ± 4.25 | 15.64 ± 4.28 | 311.11 ± 75.16<sup>c</sup> |
| Chemogenetics | Female mCherry: | 21.41 ± 8.12 | 31.30 ± 5.43 | 500.56 ± 94.11<sup>d</sup> |
|  | Male mCherry: | 13.60 ± 4.44 | 16.78 ± 7.49 | 289.89 ± 87.30<sup>d</sup> |
|  | Female hM4Di: | 20.67 ± 5.94 | 30.55 ± 6.75 | 467.89 ± 69.89<sup>e</sup> |
|  | Male hM4Di: | 13.29 ± 6.20 | 16.56 ± 5.88 | 272.53 ± 62.82<sup>e</sup> |

## Results

To determine whether a history of alcohol in rodents alters PVN^CRH^ *Fos* expression and corticosterone in early withdrawal, we subjected female and male wildtype mice to six weeks of 2-BC and brains and trunk blood was collected for FISH (*Crh* and *Fos* mRNA) and plasma corticosterone measurements (Fig 1A, M). Female water and alcohol exposed mice showed similar basal corticosterone levels (Fig 1B, unpaired two-tailed t-test: t_18_ = 0.224, p = 0.825). Interestingly, while the total number of *Fos* positive cells in the PVN was lower in alcohol exposed mice (Fig 1C, unpaired two-tailed t-test: t_17_ = 2.226, p = 0.033), the total number of *Crh* neurons (Fig 1D, unpaired two-tailed t-test: t_17_ =0.897, p = 0.382) and PVN^CRH^ activation (Fig 1E, Mann-Whitney test: [U = 28, p = 0.182]), measured as the percent of *Crh* expressing neurons that were also positive for *Fos*, was unchanged in acute withdrawal in females compared to water controls.

**Figure 1:**
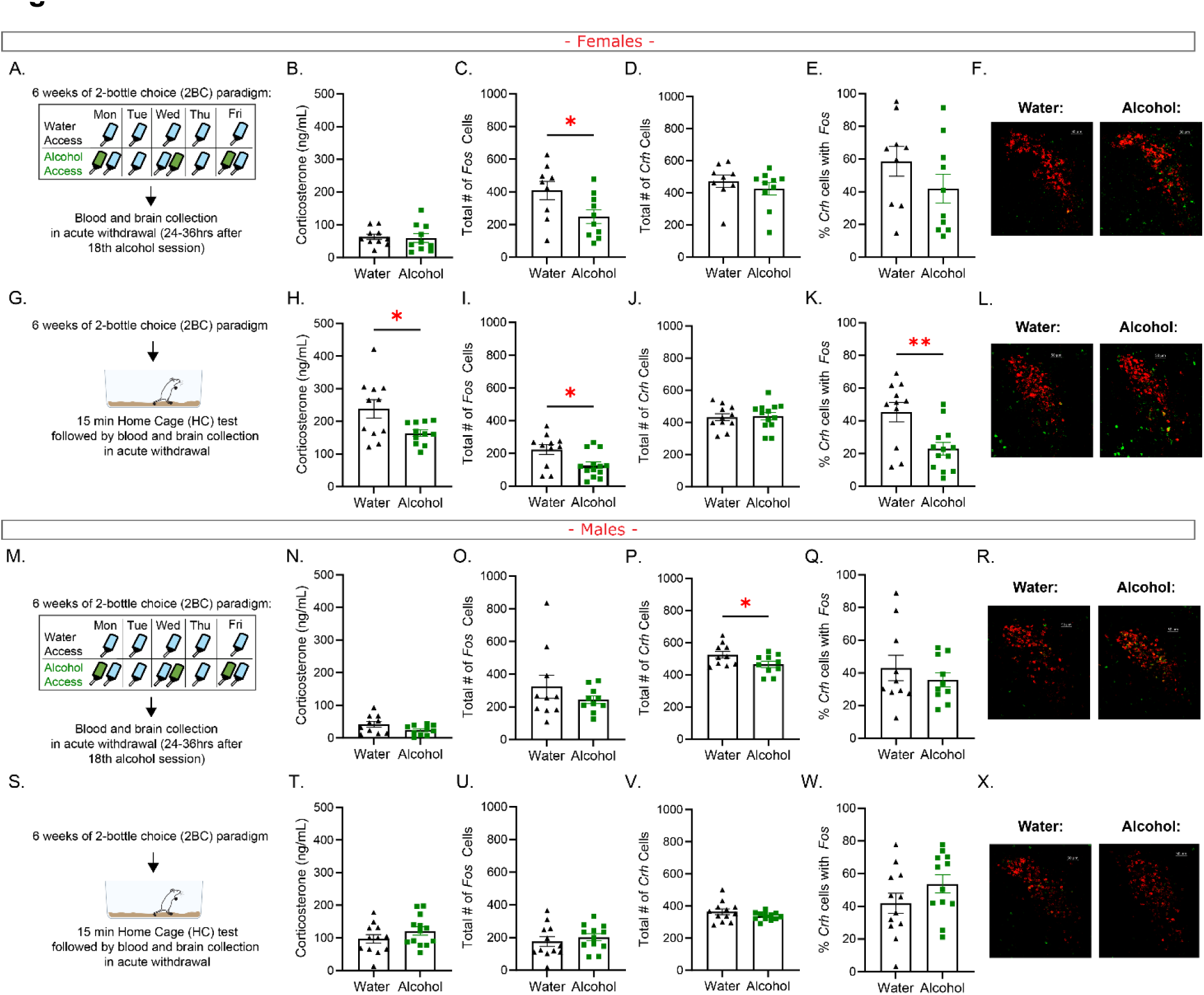
Acute stress in alcohol withdrawal blunts plasma corticosterone and PVN^CRH^ *Fos* mRNA expression in female alcohol history mice. In females: (A) experimental design for basal measurements after 2-BC alcohol (B) corticosterone (C) total number of *Fos* positive cells (D) total number of *Crh* positive cells (E) percent of *Crh* positive cells that were also positive for *Fos* (F) representative images of water (left) and alcohol (right) mice. (G) experimental design for acute stress measurements after 2-BC alcohol (H) corticosterone (I) total number of *Fos* positive cells (J) total number of *Crh* positive cells (K) percent of *Crh* positive cells that were also positive for *Fos* (L) representative images of water (left) and alcohol (right) mice. In males: (M) experimental design for basal measurements after 2-BC alcohol (N) corticosterone (O) total number of *Fos* positive cells (P) total number of *Crh* positive cells (Q) percent of *Crh* positive cells that were also positive for *Fos* (R) representative images of water (left) and alcohol (right) mice. (S) experimental design for acute stress measurements after 2-BC alcohol (T) corticosterone (U) total number of *Fos* positive cells (V) total number of *Crh* positive cells (W) percent of *Crh* positive cells that were also positive for *Fos* (X) representative images of water (left) and alcohol (right) mice.

Previous literature suggests that acute stressors can blunt PVN and hormone responses during withdrawal from vapor exposure in male rats (Lee et al., 2000). Here, tested this in a voluntary drinking paradigm by subjecting mice to six weeks of 2-BC followed by a home-cage test in acute withdrawal one day after the final alcohol session. Female mice exposed to acute stress during withdrawal had significantly lower levels of corticosterone (Fig 1H unpaired two-tailed t-test: t_20_ = 2.509, p = 0.021), fewer *Fos* positive neurons in the PVN (Fig 1H unpaired two-tailed t-test: t_22_ = 2.634, p = 0.015), and less PVN^CRH^ activation (Fig 1K Mann-Whitney two-tailed test: [U = 25, p = 0.006]) compared to water controls. This was not due to differences in *Crh* total number of neurons, as these were not different between water and alcohol exposed mice (Fig 1J unpaired two-tailed t-test: t_22_ = 0.199, p = 0.844). Male mice exposed to alcohol had similar basal corticosterone levels as water treated mice (Fig 1N, unpaired two-tailed t-test: t_18_= 1.801, p = 0.088). These two groups also had similar number of *Fos* positive PVN neurons (Fig 1O, unpaired two-tailed t-test: t_18_= 1.077, p = 0.296). While the total number of PVN *Crh* positive neurons was lower in alcohol exposed mice (Fig 1P, unpaired two-tailed t-test: t_18_= 2.108, p = 0.049), the activation of PVN^CRH^ neurons did not differ between water and alcohol exposed male mice (Fig 1E, Mann-Whitney two-tailed test: [U = 43, p = 0.631]). Acute stress during withdrawal did not affect corticosterone (Fig 1T unpaired two-tailed t-test: t_23_ = 1.312, p = 0.203), *Fos* (Fig 1U, unpaired two-tailed t-test: t_22_= 0.703, p = 0.490), total number of PVN *Crh* (Fig 1V, unpaired two-tailed t-test: t_22_= 1.427, p = 0.168) or PVN^CRH^ activation (Fig 1W, Mann-Whitney two-tailed test: [U = 49, p = 0.198]) in alcohol exposed male mice compared to water controls.

Thus far, the only electrophysiological investigations of PVN^CRH^ neurons have been conducted in protracted abstinence, long after alcohol cessation. In these experiments, we show that alcohol history also directly affects PVN^CRH^ neurons in acute withdrawal using *ex-vivo* slice electrophysiology after a home-cage test in the PVN of CRH^cre^ x Ai9 male and female mice. First, we performed voltage-and current clamp recordings to examine the intrinsic characteristics of PVN^CRH^ neurons. Female alcohol history mice had no differences in the average input resistance compared to water controls (Fig 2A, unpaired two-tailed t-test: t_34_= 0.930, p = 0.359), but did show a significantly depolarized RMP (Fig 2B: unpaired two-tailed t-test: t_34_ = 3.815, p<.001). We then measured the AP frequency in response to current steps delivered in 10pA increments from a membrane potential of -80mV. There were significant differences in intrinsic excitability between female alcohol and water mice (Fig 2C-D); a two-way RM ANOVA revealed a main effect of current (F_2.273,75.00_ = 75.59, p<.001), and a alcohol x current interaction (F_8,264_ = 2.854, p = 0.005), with no effect of alcohol (F_1,33_ = 1.466, p = 0.235). Post hoc Šidák corrected multiple comparisons test did not reveal any significant differences (30pA: p = 0.065 and 40pA: p = .108). We only found differences between alcohol and water exposed males in the input resistance (Fig 2E unpaired two-tailed t-test: t_30_ = 2.263, p = 0.031), with no differences in the RMP (Fig 2F: unpaired two-tailed t-test: t_30_ = 0.890, p = 0.381), or current step protocol (Fig 2G-H).

**Figure 2:**
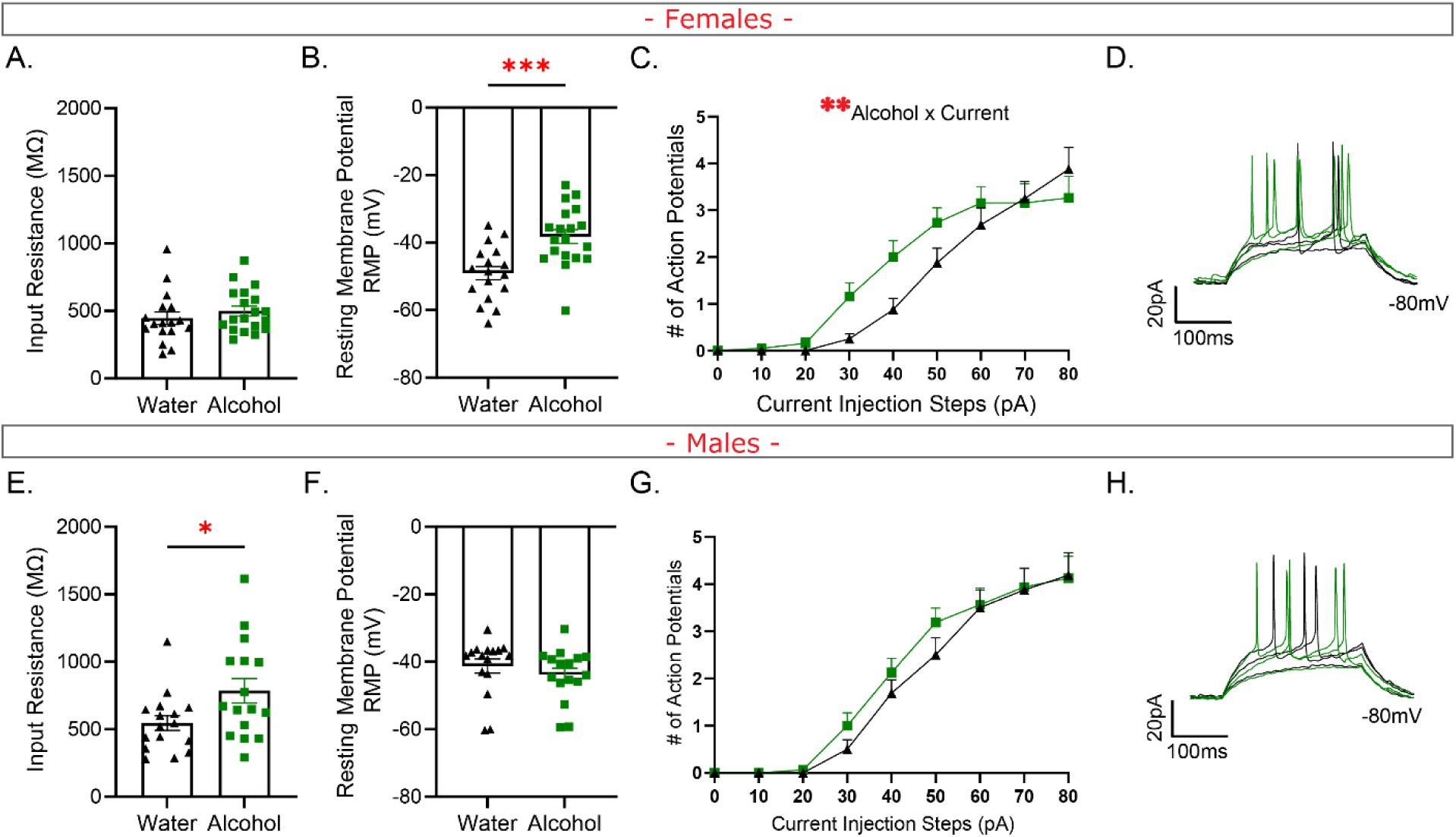
Chronic alcohol consumption engenders a hyperexcitable phenotype in PVNCRH neurons in female alcohol withdrawal mice. In females: (A) Input resistance (B) Resting membrane potential (RMP) in mV (C) Current step injection plot (D) Representative 30 – 50pA current steps in current step injection plot. In males: (E) Input resistance (F) Resting membrane potential (RMP) in mV (G) Current step injection plot (H) Representative 30 – 50pA current steps in current step injection plot. For A-B, E-F: Student unpaired t-test. For C & E: two-way repeated measures ANOVA. * p < .05, ** p < .01, *** p < .001.

Having established differences in intrinsic excitability in females, but not males, we next measured synaptic transmission in PVN^CRH^ neurons (Fig 3). As hypothesized, female alcohol history mice had stronger excitatory synaptic drive compared to water mice, as mEPSC frequency was higher in these mice (Fig 3A unpaired t-test: t_13_ = 2.426, p = 0.025) and sEPSC frequency was trending higher in alcohol mice (Fig 3D unpaired t-test: t_29_ = 1.944, p = 0.062). There was no difference in the amplitude of mEPSCs (Fig 3B) or sEPSCs (Fig 3E) between female mice. Female alcohol and water mice also had no differences in mIPSC frequency (Fig 3G) and amplitude (Fig 3H) or sIPSC frequency (Fig 3J) and amplitude (Fig 3K). Interestingly, male alcohol history mice also displayed a high frequency of mEPSCs compared to water controls (Fig 3M unpaired t-test: t_22_ = 2.970, p = 0.007) with no difference in mEPSC amplitude (Fig 3N) or sEPSCs frequency (Fig 3P) or amplitude (Fig 3Q). However, there were also increases in inhibitory synaptic transmission in alcohol history male mice compared to water controls. While there were no changes in mIPSC frequency (Fig 3S), there was a significant increase in mIPSC amplitude (Fig 3T unpaired t-test: t_24_ = 2.348, p = 0.028), with sIPSCs showing increased frequency (Fig 3V unpaired t-test: t_30_ = 2.484, p = 0.019) but no differences in amplitude (Fig 3W).

**Figure 3:**
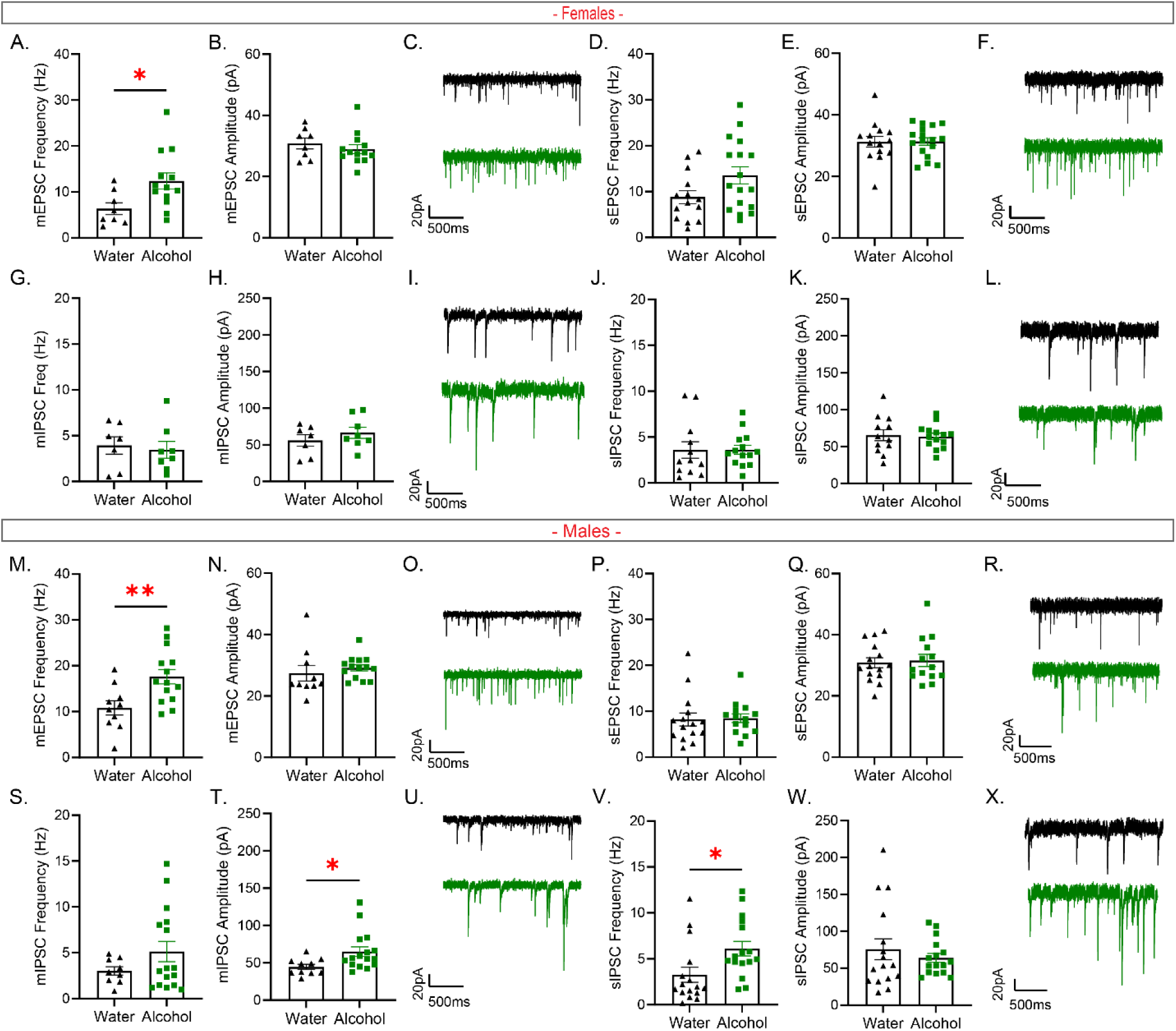
A history of alcohol increases glutamatergic synaptic transmission in female and male alcohol withdrawal mice and GABAergic synaptic transmission in male alcohol withdrawal mice. In females: (A) miniature excitatory post synaptic current (mEPSC) frequency (B) mEPSC amplitude (C) representative traces for water (black) and alcohol (green) mEPSCs (D) spontaneous excitatory post synaptic current (sEPSCs) frequency (E) sEPSC amplitude (F) sEPSC amplitude (F) representative traces for water (black) and alcohol (green) sEPSCs (G) miniature inhibitory post synaptic current (mIPSC) frequency (H) mIPSC amplitude (I) representative traces for water (black) and alcohol (green) mIPSCs (J) spontaneous inhibitory post synaptic current (sIPSC) frequency (K) sIPSC amplitude (L) representative traces for water (black) and alcohol (green) sIPSCs. In males: (M) mEPSC frequency (N) mEPSC amplitude (O) representative traces for water (black) and alcohol (green) mEPSCs (P) sEPSC frequency (Q) sEPSC amplitude (R) representative traces for water (black) and alcohol (green) sEPSCs (S) mIPSC frequency (T) mIPSC amplitude (U) representative traces for water (black) and alcohol (green) mIPSCs (V) sIPSC frequency (W) sIPSC amplitude (X) representative traces for water (black) and alcohol (green) sIPSCs. Student unpaired t-tests. * p < .05, ** p < .01.

Finally, we tested whether in-vivo chemogenetic inhibition of PVN^CRH^ neurons during withdrawal can alter behavior. Virus placements are represented in Fig 4B. First, 24-hrs after the final alcohol session, mice were subjected to the home-cage test immediately followed by a free social interaction test (Fig 4A). Female mice, regardless of alcohol history or hM4Di mediated inhibition of PVN^CRH^ neurons, displayed similar behavioral phenotypes in the home cage test (Fig 4C). Using a two-way ANOVA, we did not find any significant main effects or interaction effect of alcohol or hM4Di in the distance moved (Fig 4D), time spent rearing (Fig 4E), time spent digging (Fig 4F) or time spent grooming (Fig 4G). Female alcohol and water exposed mice also displayed no differences in the social interaction test (Fig 4H), as determined by two-way ANOVAs, in face contact (Fig 4I), anogenital contact (Fig 4J main effect of hM4Di F_1,42_ = 3.439, p = 0.071), allogrooming (Fig 4K main effect of hM4Di F_1,42_ = 3.267, p = 0.078) or fighting (Fig 4L main effect of hM4Di F_1,42_ = 2.928, p = 0.094). Male hM4di mice in the home-cage test had significantly reduced distance moved (Fig 4N, two-way ANOVA main effect of hM4Di F_1,38_ = 4.546, p = 0.040) regardless of alcohol history. Time spent rearing (Fig 4O), digging (Fig 4P), and grooming (Fig 4Q) was not affected by alcohol history or virus. Interestingly, alcohol history had a significant effect on the social interaction test in males (Fig 4R). Alcohol history mice, regardless of virus, spent more time in face (Fig 4S two-way ANOVA main effect of alcohol F_1,37_ = 6.150, p = 0.018) and anogenital contact (Fig 4T two-way ANOVA main effect of alcohol F_1,37_ = 14.04, p < .001) with a juvenile same sex mice than water controls. There was also a trend towards an effect of hM4Di on reduced anogenital contact (two-way ANOVA main effect of hM4Di F_1,37_ = 3.899, p = 0.056). Allogrooming (Fig 4U) and time spent fighting was not affected by alcohol or virus in male mice (Fig 4V).

**Figure 4:**
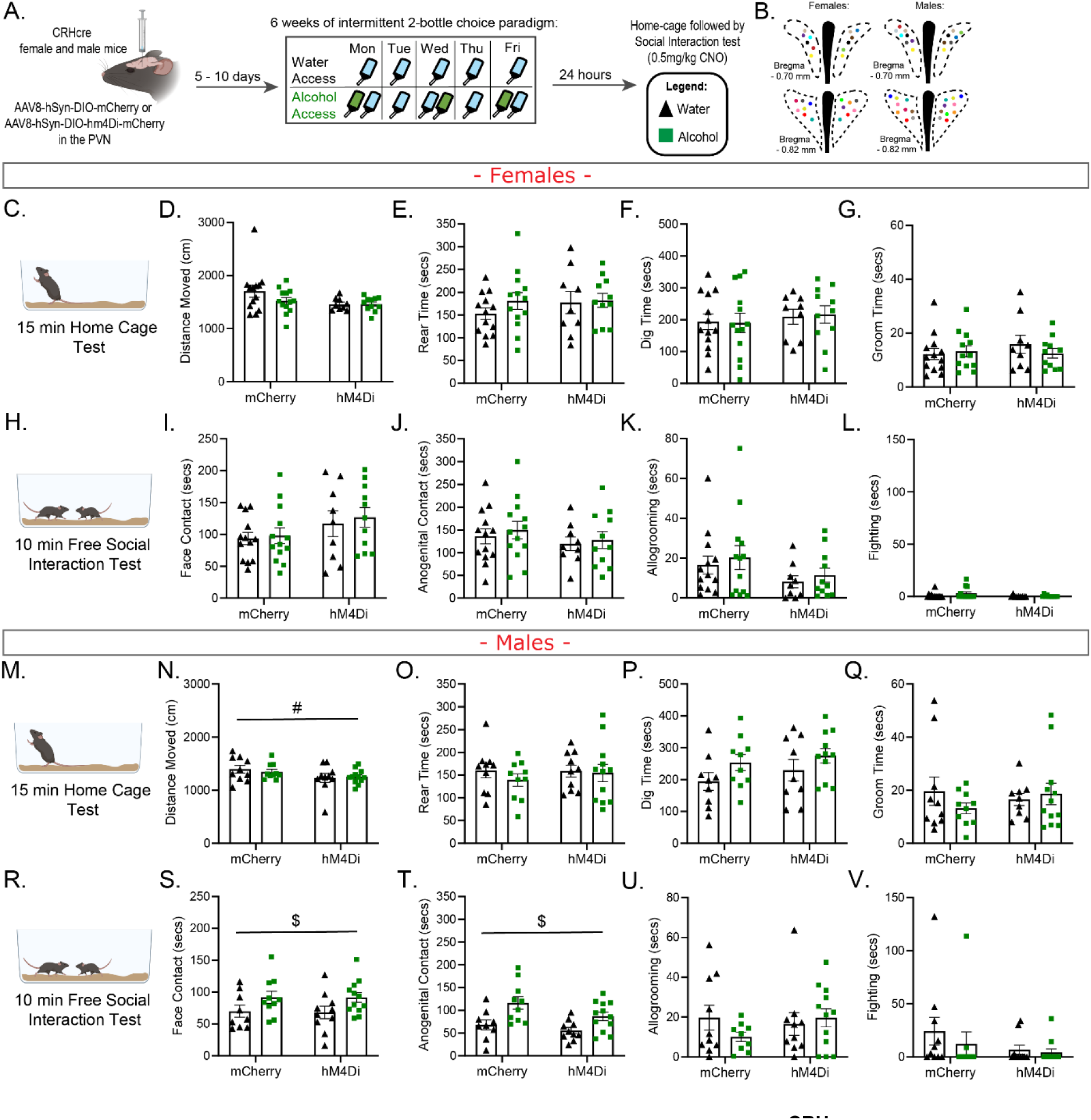
hM4Di mediated chemogenetic inhibition of PVN^CRH^ neurons did not affect mouse responses to the home-cage or free social interaction test in acute withdrawal from chronic alcohol. (A) Experimental design for female and male CRH-cre mice. (B) Virus placements, each color represents one mouse. In females: (C) 15- min home-cage test (D) Distance moved (E) Rear time (F) Dig time (G) Groom time (H). 10-min free social interaction test (I) Face contact (J) Anogenital contact (K) Allogrooming (L) Fighting. In males: (M) 15-min home-cage test (N) Distance moved (O) Rear time (P) Dig time (Q) Groom time. (R) 10-min free social interaction test (S) Face contact (T) Anogenital contact (U) Allogrooming (V) Fighting. Two-way repeated measures ANOVA. #: Main effect of hM4Di virus, $: Main effect of alcohol history

We had previously shown (Neira et al., 2022) that female mice in 6-8hr acute withdrawal had increased escape behaviors in the looming disc test, therefore, we hypothesized that inhibiting PVN^CRH^ neurons will restore female behavior in this test. First, we established that increased escape responses compared to mCherry water controls in 6- day alcohol withdrawal mCherry mice were also seen in the looming disc test (Fig 5A).

**Figure 5:**
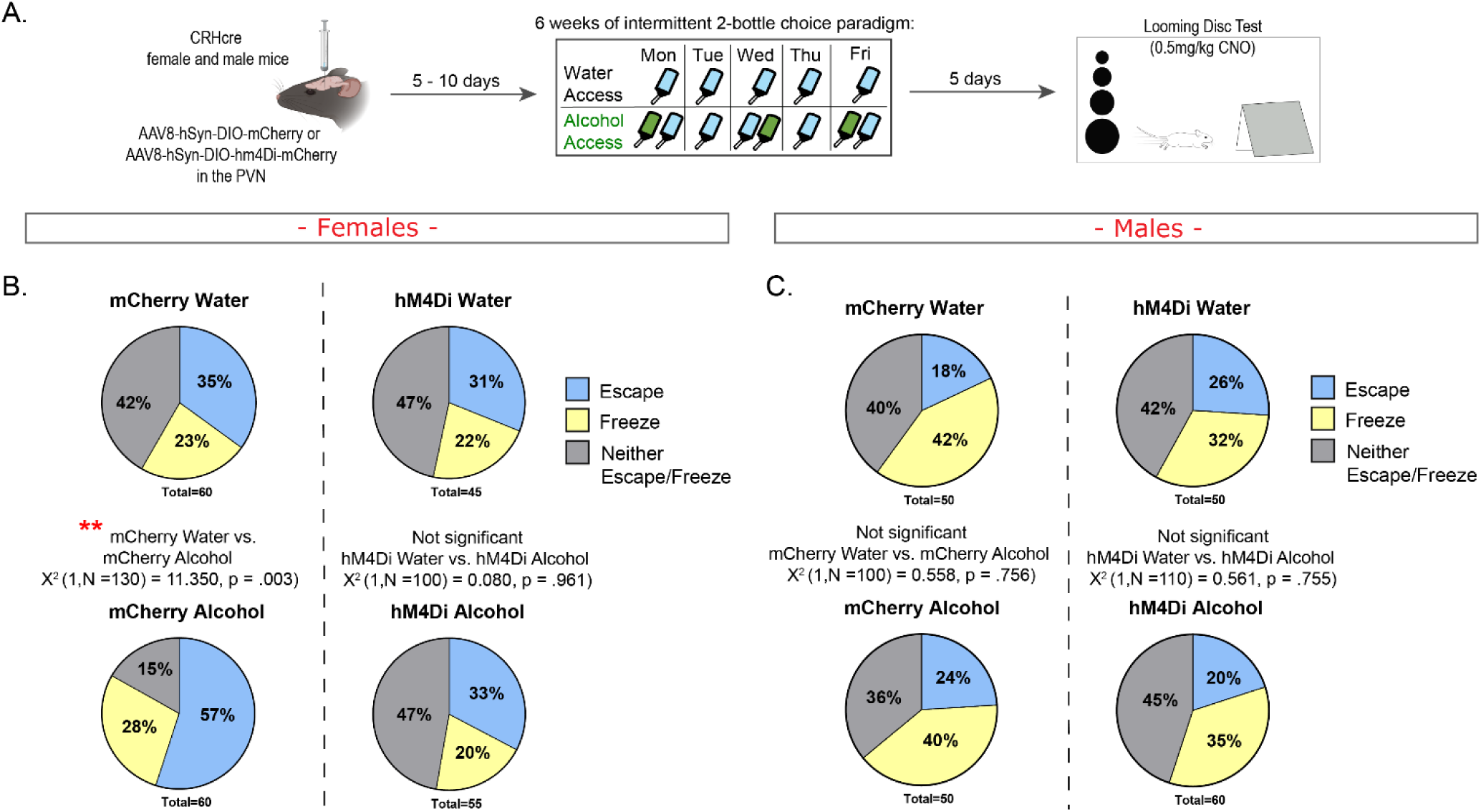
hM4Di mediated chemogenetic inhibition of PVN^CRH^ neurons restores looming disc response behaviors in female alcohol history mice during withdrawal. (A) experimental design (B) Female escape (blue), freeze (yellow) or neither escape or freeze (grey) response to loom presentation (C) Male escape (blue), freeze (yellow) or neither escape or freeze (grey) response to loom presentation. Chi- square. ** p < .01.

In fact, Chi-square analysis between mCherry water and alcohol mice during Escape, Freeze, or neither escape/freeze behavioral responses to loom revealed significant differences in behavior (Fig 5B X^2^ (1, N =130) = 11.3502, p =.003), with alcohol mCherry mice displaying greater escape responses. Importantly, hM4Di water and alcohol history female mice had near identical reactions to the looming disc test, as indicated by the non-significant Chi-square test (Fig 5B X^2^ (1, N =100) = 0.080, p =.961). Unlike females, male mCherry alcohol history mice and water controls did not differ in their response to loom presentations (Fig 5C), similar to what we had previously shown (Neira et al., 2022). Importantly, hM4Di water and alcohol history male mice also showed similar responses to the looming disc task (Fig 5C).

## Discussion

In this study, we find that a history of voluntary alcohol consumption alters PVN^CRH^ neurons during acute withdrawal and that inhibition of these neurons restores aberrant stress coping behaviors in female mice. Specifically, using FISH, we find that mRNA *Fos* in PVN^CRH^ neurons is unaltered in basal conditions (Fig 1E) but blunted in response to an acute stressor in female mice (Fig 1K). This is similar to what has been shown in humans AUD patients and in male rats using forced vapor experiments. Nonetheless, male mice in this voluntary drinking paradigm did not show any alterations to *Fos* in PVN^CRH^ neurons in basal (Fig 3Q) or stress conditions (Fig 3W), perhaps due to lower levels of alcohol intake in males. Similar to these FISH findings, plasma corticosterone levels were only blunted in female alcohol history mice exposed to an acute stressor (Fig 1H).

Despite *Fos* mRNA indicating a reduction in the activation of PVN^CRH^ neurons during acute withdrawal, slice electrophysiology experiments demonstrated PVN^CRH^ are in a hyperactive state, as measured through intrinsic excitability and synaptic transmission experiments (Fig 2 & 3). These effects were more prominent in female mice.

Interestingly, males showed an increase in inhibitory synaptic transmission (Fig 3T & V) that may act as a protective mechanism against alcohol induced increases in glutamatergic signaling during acute withdrawal. *Fos* alterations are often conceptualized as changes to membrane depolarization, however studies have found *Fos* induction can also be a response to intracellular signaling mechanisms, and as such does not always correlate with neuronal depolarization, making it a blunt tool for interpreting neuronal activation (Hoffman et al., 1993; Hoffman & Lyo, 2002; Joo et al., 2016). Slice electrophysiology is a more direct tool that can be used to measure the activity of a neuron, and in these studies, we find that *Fos* and slice electrophysiology experiments findings are mismatched. This might be due to an uncoupling of the mechanisms for neuroendocrine HPA-axis activation and central PVN^CRH^ neuronal activity, as *Fos* induction has been thought to invoke peptide release and has been used to study neuroendocrine function (Hoffman & Lyo, 2002), or could be indicative of distinct populations of PVN^CRH^ neurons with neuroendocrine and central functions (Bains et al., 2015; Lameu et al., 2022). Our plasma corticosterone and *Fos* activation studies align, and perhaps PVN^CRH^ neurons regulate CRH release into the periphery differently than glutamate release centrally. In fact, many recent studies have found that these neurons are involved in various stress related behaviors independent of their neuroendocrine function (Daviu et al., 2020; Füzesi et al., 2016; Kim et al., 2019; Yuan et al., 2019).

Female, not male, alcohol history mice in acute 6-8 hour withdrawal show greater active responses to the looming disc test when compared to water controls (Neira et al., 2022). In line with the findings indicating increased intrinsic activity and glutamatergic function in alcohol history female mice, we hypothesized that these aberrant behaviors seen in acute withdrawal female mice in the looming disc may be due to increased activity of PVN^CRH^ neurons, as inhibition of these neurons reduces escape behaviors in this test (Daviu et al., 2020). In this study, we further confirm increased active response in the looming disc task in female, not male, alcohol history mice in 5-day withdrawal.

Importantly, we also show that in-vivo chemogenetic inhibition of PVN^CRH^ neurons restores female alcohol history mice behavior to the looming disc (Fig 5). Overall, alcohol induced hyperactivity of PVN^CRH^ neurons drives heightened active stress coping in female mice in acute withdrawal, and inhibition of these neurons is sufficient to restore these stress coping behaviors.

### Plasma Corticosterone and FISH in the PVN

To our knowledge, this is the first study to report *Fos* induction in PVN^CRH^ neurons after voluntary drinking in male and female rodents. Of interest, basal levels of *Fos* activation in PVN^CRH^ neurons were highly varied in both females and males. For our basal condition experiments, mice were removed from their animal facility housing room after anesthesia, as we tried to be as close to a no-stress condition as possible. It was intriguing to note that despite these measures, some mice had 80-100% *Fos* activation in *Crh* neurons. This variability was mostly also present in stress conditions for males and water female mice, while alcohol female mice did not appear to mount a significant stress response, which aligned with their blunted plasma corticosterone levels. Similar findings with blunting of c-fos response are seen in rodents after chronic stress conditions (Armario et al., 2004; Borrow et al., 2019; Walker et al., 2019). A few studies have also found reduced hormonal and c-fos response to stress after ethanol pre-exposure (Lee et al., 2000; Lee & Rivier, 1993; Rivier et al., 1990). These mice were single housed for extended periods, which is itself a chronic stressor that can affect PVN^CRH^ neurons, particularly in females. It is possible that chronic alcohol, withdrawal episodes and social isolation all contributed to a chronic stress phenotype in alcohol history mice. Experiments to delineate the contributions of each of these will be important to further understand alcohol induced changes in stress function.

### Slice ex-vivo Electrophysiology in PVN^CRH^ neurons

Many studies have shown that chronic stress induces a hyperexcitable state in PVN^CRH^ neurons (Flak et al., 2009; Franco et al., 2016; Stephens & Wand, 2012). Acute alcohol acts as a stressor and induces the HPA-axis system, and withdrawal itself is also a stressor on the body. Thus, chronic alcohol consumption with repeated withdrawals is likely acting as a chronic stressor and inducing a hyperactive condition in PVN^CRH^ neurons. Our electrophysiological data supports this hypothesis, particularly in female mice, as they display both increased intrinsic excitability (Fig 2) and increased glutamatergic synaptic transmission (Fig 3). Interestingly, male alcohol history mice also shown increased glutamatergic transmission, but do not display increased intrinsic excitability apart from an increase in input resistance (Fig 2). Males drink less alcohol in 2-BC procedures than females (Neira et al., 2022), which might account for the lack of some effects and justifies an investigation with alcohol vapor to normalize total alcohol levels. Alternatively, males may have a mechanism of protection against alterations to intrinsic excitability. In fact, males show an increase in inhibitory synaptic transmission, both in the frequency of sIPSCs and amplitude of mIPSCs (Fig 3), perhaps indicating global increases in inhibitory inputs that allow for homeostasis along with potential post synaptic adaptations. Notably, we did observe some differences between measures of spontaneous and miniature transmission in our experiments. These could be due to multiple reasons. For example, the difference in spontaneous and miniature EPSC frequency in males could be due to alterations in action potential dependent release of other signaling molecules, such as peptides. The difference in spontaneous and miniature IPSC amplitude in males could potentially be driven by engagement of different pools of vesicles across condition or perhaps driven by other tonic currents.

Overall, females might be more sensitive to PVN^CRH^ alterations from alcohol and thus suffer from the consequences of heighted activity of these neurons. If females are more affected by alcohol changes to PVN^CRH^ neurons, it might partially explain the differences in reported reasons for drinking between men and women, as women often report drinking to alleviate stressors.

### hM4Di Chemogenetic Inhibition of PVN^CRH^ neurons

In support of the hypothesis that females are more sensitive to PVN^CRH^ alterations from alcohol, we show that female alcohol history mice display greater active response behaviors in the looming disc test, and that chemogenetic inhibition restores escape behaviors in hM4Di female mice, as alcohol and water mice behavior is indistinguishable (Fig 5). This indicates that altered behaviors in response to loom presentation in alcohol history female mice are tied to increased PVN^CRH^ activity.

Manipulations of PVN^CRH^ neurons also alter spontaneous behaviors such as rearing. Rearing is significantly increased during optogenetic inhibition of PVN^CRH^ neurons following a stressful event and reduced during optogenetic activation in different contexts (Füzesi et al., 2016). Commonly thought of as exploratory (Sturman et al., 2018), time spent rearing appears to be sensitive to context, as it is commonly seen in home-cage and novel environments, but less common in a high stress environment, such as the foot shock chambers immediately after a shock stress (Füzesi et al., 2016; Neira et al., 2022). Grooming behaviors have also been bi-directionally driven by activation and inhibition of PVN^CRH^ neurons (Füzesi et al., 2016; Yuan et al., 2019).

Thus, we expected to find alterations to spontaneous behaviors such as rearing and grooming in the home-cage during chemogenetic inhibition of PVN^CRH^ neurons, as has also been seen in acute withdrawal from chronic alcohol (Neira et al., 2022). In contrast, we did not see any changes to these behaviors in a home-cage test during acute withdrawal in female or male hM4Di mice (Fig 4). This might be due to the relatively low dose of CNO that was used in these studies that perhaps did not override natural behavior patterns like optogenetics. We also failed to see any alcohol history driven effects in the home-cage test, which might be due to the stress of the CNO injection given before the test. In fact, the behaviors displayed by the mice were similar to those displayed by acute restraint stress mice in our previously published manuscript (Neira et al., 2022). Immediately after the home-cage test, we conducted a free social interaction test. Interestingly, while we saw no effects of PVN^CRH^ inhibition on female or male mice, male alcohol history mice spent significantly more time interacting with a juvenile mouse that water controls. Literature suggests that females are more likely to suffer from alcohol induced reductions in sociability (Flanigan et al., 2022; Sidhu et al., 2018; Simon et al., 2023), however, this is the first study to our knowledge that has investigated free social interaction in acute withdrawal from voluntary two-bottle choice alcohol consumption in mice.

Because manipulations of PVN^CRH^ neurons might alter behavior in context and dose dependent manners, follow up experiments using higher doses of CNO, chemogenetically activating these neurons, and using optogenetics will be highly informative to understand the role PVN^CRH^ neurons play in alcohol induced stress behaviors. It will also be imperative, through single-cell imaging experiment, to understand the natural activity of PVN^CRH^ neurons in response to acute alcohol sessions, as well as during different stress response tests.

## Conclusions

Overall, as summarized in our working model in Figure 6, alcohol history mice demonstrate significant alterations to PVN^CRH^ neurons in withdrawal, with females indicating more expansive alterations to these neurons and stress related behaviors that involve the activity of these neurons. Perhaps this is due to the nature of voluntary intake in this study which leads to lower alcohol consumption in males than females, or to the social isolation that is required for 2-BC experiments that affects females greatly (Cacioppo et al., 2011; Senst et al., 2016). Alternatively, females, who already have a more active stress system in response to multiple stressors (Bangasser et al., 2018; Palanza & Parmigiani, 2017; Peltier et al., 2019; Silva et al., 2009), could be more sensitive to alcohol induced changes in PVN^CRH^ neurons. These changes might further impact their ability to cope with stressors, which might play a role in reports of women drinking for stress relief rather than the rewarding effects of alcohol consumption (Pollard et al., 2020; Rodriguez et al., 2020).

**Figure 6:**
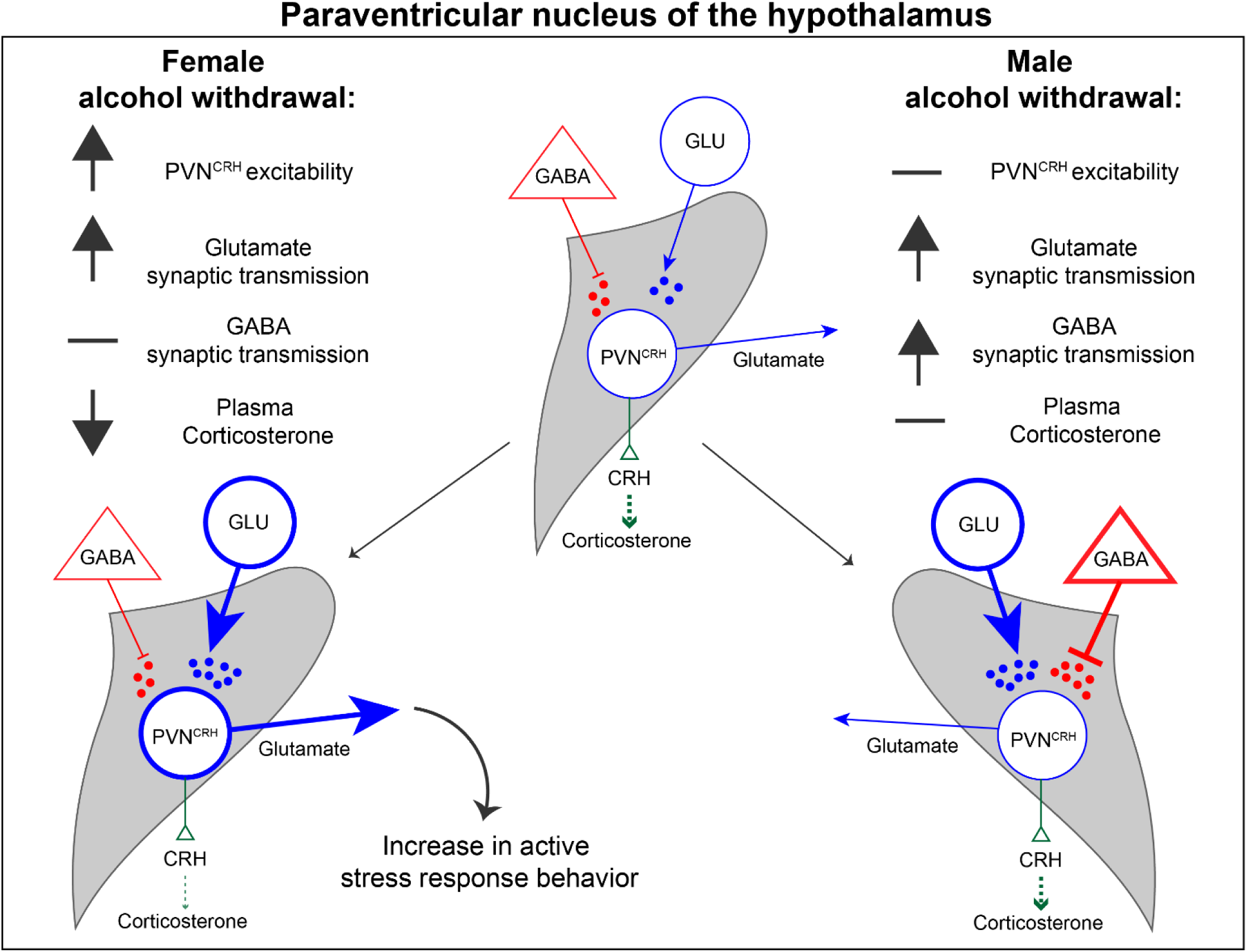
Working model of paraventricular nucleus of the hypothalamus corticotropin releasing hormone (PVN^CRH^) neuron alterations in female and male mice. Summary of findings in females (left) and males (right) in withdrawal from a chronic history of alcohol consumption.

